# Genomic and TCR Repertoire Intratumor Heterogeneity of Small-cell Lung Cancer and its Impact on Survival

**DOI:** 10.1101/2020.06.30.180844

**Authors:** Ming Chen, Runzhe Chen, Ying Jin, Jun Li, Jiexin Zhang, Junya Fujimoto, Won-Chul Lee, Xin Hu, Shawna Maria Hubert, Julie George, Xiao Hu, Yamei Chen, Carmen Behrens, Chi-Wan Chow, Hoa H.N. Pham, Junya Fukuoka, Edwin Roger Parra, Carl M. Gay, Latasha D. Little, Curtis Gumbs, Xingzhi Song, Lixia Diao, Qi Wang, Robert Cardnell, Jianhua Zhang, Jing Wang, Don L. Gibbons, John V. Heymach, J. Jack Lee, William N. William, Bonnie Glisson, Ignacio Wistuba, P. Andrew Futreal, Roman K. Thomas, Alexandre Reuben, Lauren A. Byers, Jianjun Zhang

## Abstract

Small-cell lung cancer (SCLC) is speculated to harbor complex genomic intratumor heterogeneity (ITH) associated with high recurrence rate and suboptimal response to immunotherapy. Here, we revealed a rather homogeneous mutational landscape but extremely suppressed and heterogeneous T cell receptor (TCR) repertoire in SCLCs. Higher mutational burden, lower chromosomal copy number aberration (CNA) burden, less CNA ITH and less TCR ITH were associated with longer overall survival of SCLC patients. Compared to non-small cell lung cancers (NSCLCs), SCLCs had similar predicted neoantigen burden and mutational ITH, but significantly more suppressed and heterogeneous TCR repertoire that may be associated with higher CNA burden and CNA ITH in SCLC. Novel therapeutic strategies targeting CNA could potentially improve the tumor immune microenvironment and response to immunotherapy in SCLC.

## Introduction

Small-cell lung cancer (SCLC) accounts for ∼15% of all newly diagnosed lung cancers leading to ∼30,000 deaths in the United States annually^1^. SCLC is a highly aggressive cancer characterized by rapid growth and high rates of early local and distant metastases^2,3^. At initial diagnosis, around one third of SCLC patients present with cancer confined to one hemithorax, defined as limited-stage disease (LD) that can be treated with chemotherapy combined with radiotherapy or surgical resection, while the remaining patients present with extensive-stage disease (ED) exhibiting extensive lymph node involvement and/or distant metastases usually treated with palliative chemotherapy with or without immune checkpoint blockade (ICB)^4-6^. Although most SCLC patients experience an initial response, nearly all patients recur with rapidly progressing disease resistant to late-line treatments. Despite extensive research, only modest advances have been achieved in the treatment of SCLC over the past 30 years with median survival less than a year and 5-year overall survival (OS) is below 7% for ED SCLC^1,7-9^. Recently, the addition of ICB to chemotherapy has become a new standard of care for advanced SCLC, although it confers only an improvement of 2-3 months in survival^8^. National Cancer Institute (NCI) has identified SCLC as a recalcitrant malignancy^1^. Translational studies to understand the mechanisms underlying recurrence and therapeutic resistance remain an unmet need to design novel therapeutic strategies^8,10,11^.

Tumors are composed of cancer cells and stromal cells of distinct molecular and phenotypic features, a phenomenon termed intratumor heterogeneity (ITH). ITH has been shown to impact response to therapy and patient survival^12-15^. We and others have previously delineated the ITH architecture of non-small cell lung cancers (NSCLCs) at genomic, epigenetic and gene expression levels utilizing multiregional sequencing and demonstrated that complex ITH was associated with inferior survival^16-22^. It has been speculated that SCLC has extremely complex ITH architecture that lead to poor prognosis^23^. Another plausible explanation for the poor outcome is that SCLC is associated with an immunosuppressive tumor microenvironment, particularly T cell responses^24,25^. In localized NSCLC, our recent work has revealed that a suppressed and heterogeneous T cell receptor (TCR) repertoire is associated with inferior survival^18,26^.

The genomic and TCR ITH architecture of SCLC and their potential clinical impact have not been well studied, largely due to lack of available tumor specimens^27-30^. Through international collaboration, we conducted multiregional whole exome sequencing (WES) and TCR sequencing of 50 tumor samples from 19 resected LD SCLCs **(Figure S1)** to depict the immunogenomic ITH architecture of SCLC. We further compared these SCLCs to a cohort of 216 localized NSCLCs (PROSPECT cohort) ^26^ and assessed the impact of genomic and TCR attributes on patient survival.

## Results

### Mutational landscape of LD SCLC tumors is overall homogeneous

A total of 50 tumor regions (hereafter referred as region) from 19 resected LD SCLC tumors (hereafter referred as tumor) were subjected to WES (**Figure S1, Supplementary Table 1**). In total, 3,773 nonsilent (nonsynonymous, stop-gain and stop-loss) mutations were identified from these 50 tumor regions (**Supplementary Data 1**) for a median nonsilent tumor mutational burden (TMB) of 4.69/Mb. TMB was similar between different tumor regions within the same tumors, but varied substantially between patients (**Figure S2**).

Next we constructed phylogenetic trees of 18 SCLCs for which multiregional WES data available (P13 only had one tumor region and was excluded from this analysis) to depict the genomic ITH and the evolutionary trajectory of these SCLCs as described previously^17^. A median of 80.4% (28%-93%) of mutations were mapped to the trunks of these 18 SCLCs (**Figure 1** and **Figure 2A**) representing ubiquitous mutations present in all tumor regions within the same tumor, comparing to 73% (8%-99.6%) trunk mutations in 100 early-stage NSCLC in TRACERx cohort (p=0.218)^19^. Furthermore, using PyClone^31^, we classified mutations as clonal (defined as estimated cancer cell fraction=1, indicating mutations present in all cancer cells) or subclonal (defined as estimated cancer cell fraction<1, indicating mutations only present in a subset of cancer cells) in each tumor specimen. A median of 92.8% (37%-99.9%) of mutations were clonal in these 50 SCLC specimens (**Figure 2B**). Taken together, these results suggested the mutational landscape is overall homogeneous in these SCLCs.

**Figure 1.**
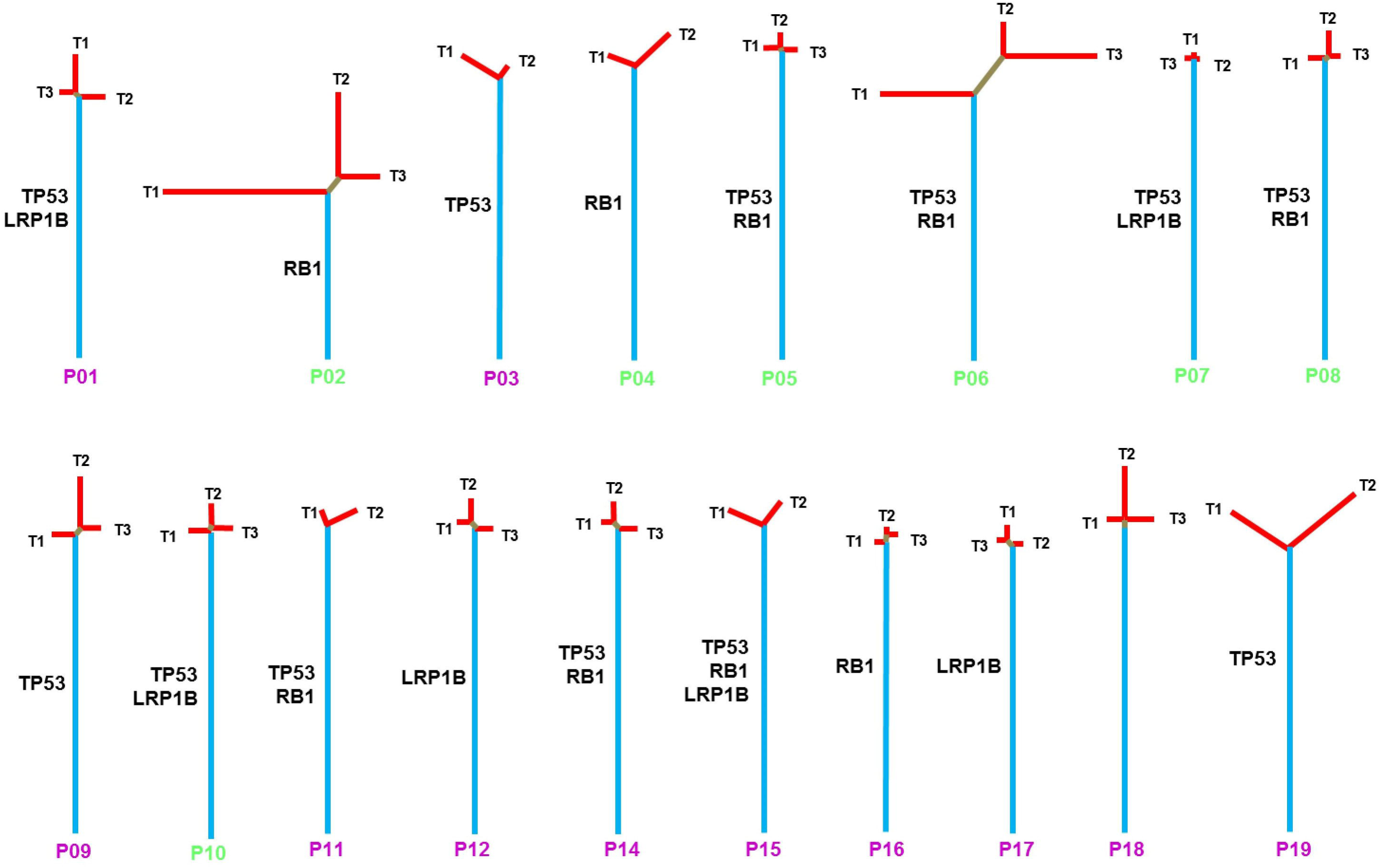
Phylogenetic trees of 18 SCLC tumors with multiregional whole exome sequencing (WES). Blue, brown and red lines represent trunk, branch, and private mutations, respectively. The length of trunk, branch and private branch are proportional to the numbers of mutations. Commonly mutated cancer genes *TP53, RB1* and are mapped to the phylogenetic trees as indicated. Patient ID: pink = alive; green = expired.

**Figure 2.**
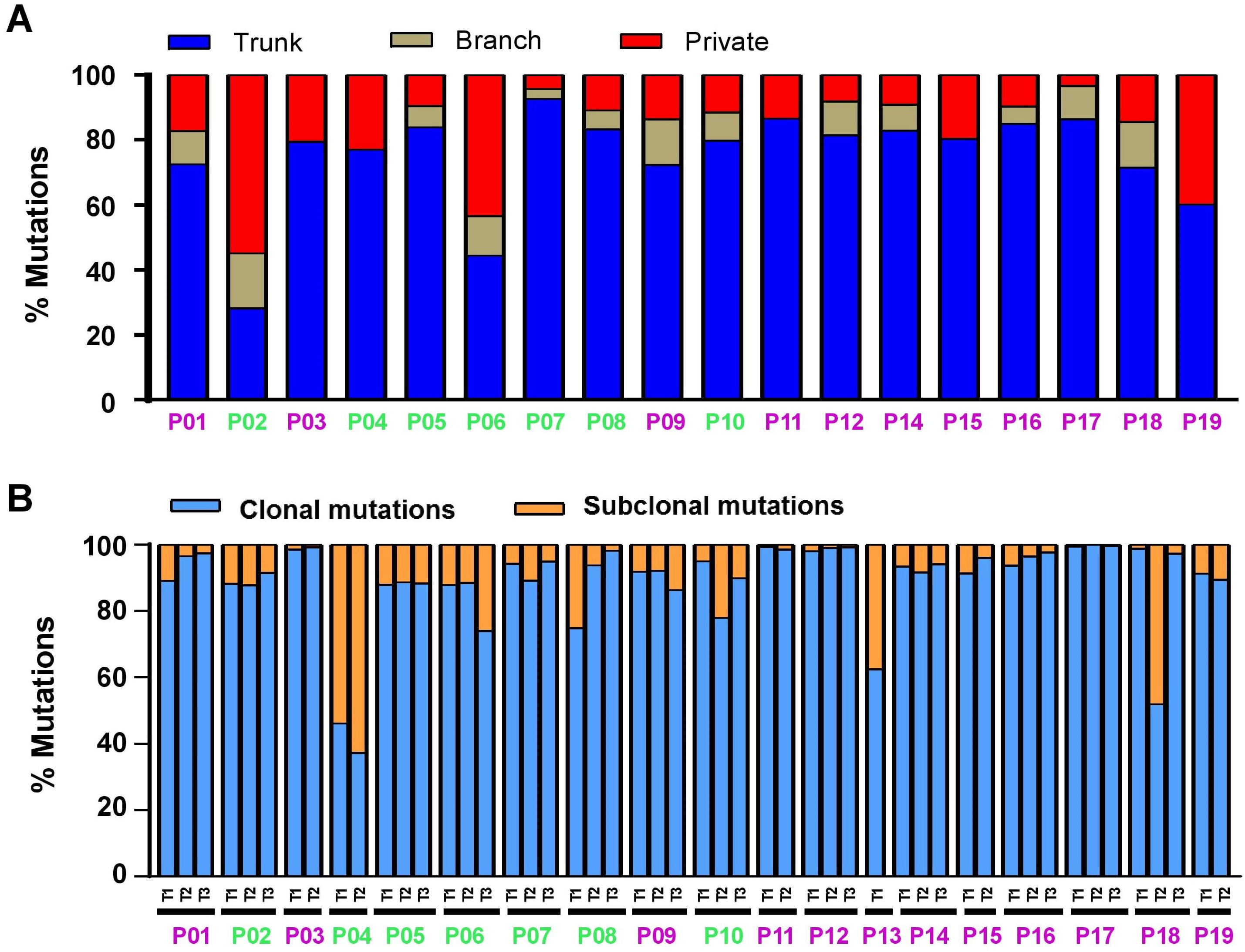
Genomic intra-tumor heterogeneity of small cell lung cancers. **(A)** Proportion of trunk (blue), branch (brown), and private (red) mutations representing mutations detected in all tumor regions, some but not all and only in one single tumor region from any given tumor. Purple patient IDs = patients who were alive; Green patient IDs = patients who were decreased. **(B)** Proportion of clonal *versus* subclonal mutations defined by PyClone in 50 SCLC tumor specimens. Mutations were classified as clonal (estimated cancer cell fraction = 1, indicating mutations presenting in all cancer cells, blue) or subclonal (estimated cancer cell fraction < 1, indicating mutations only present in a subset of cancer cells, orange) in each tumor specimen.

The most frequently mutated cancer genes in this cohort included *TP53, RB1*, and *LRP1B* identified in 14, 10 and 7 patients respectively (**Figure S3**). Of note, these mutations were trunk mutations detected in all regions from the same SCLC tumors (**Figure 1** and **Figure S3**) and these canonical mutations were clonal in each tumor specimen where they were identified. These results suggest that these canonical cancer gene mutations were all early genomic events during evolution of SCLC.

### Mutational processes in this cohort of SCLC

Understanding how mutational processes shape cancer evolution may inform mechanisms underlying tumor adaptation. We next analyzed the mutational spectrum and signatures in these SCLCs^32^. C>A transversions were the most common nucleotide substitutions (**Figure S4**) and Cosmic Signature 4 (associated with cigarette smoking) was the predominant mutational signature (**Figure 3A**) as expected, given 15 of the 19 patients were smokers.

**Figure 3.**
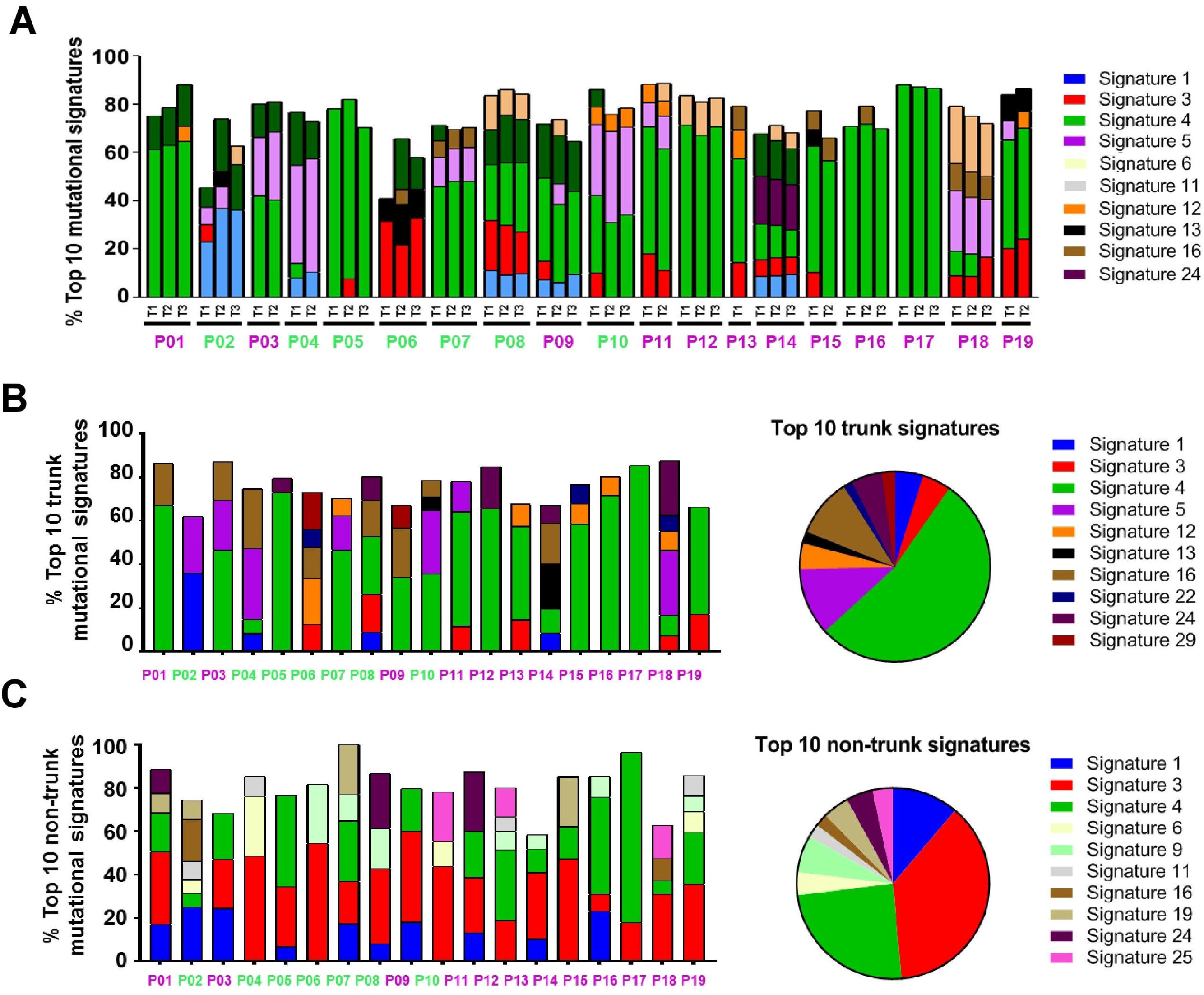
The mutational processes in small cell lung cancers. **(A)** The top COSMIC mutational signatures in 50 SCLC specimens. **(B)** The top COSMIC mutational signatures associated with trunk mutations. Bar chart on the left: top signatures associated with trunk mutations in each patient. Pie chart on the right: the average of contribution of each signature across the 19 patients. **(C)** The top COSMIC mutational signatures associated with non-trunk mutations. Bar chart on the left: top signatures associated with non-trunk mutations in each patient. Pie chart on the right: the average of contribution of top signatures across the 19 patients. Patient ID: pink = alive; green = expired.

To further dissect the mutational processes associated with early clonal expansion versus subsequent subclonal diversification, we delineated the mutational signatures of trunk mutations representing early genomic events and non-trunk mutations representing later subclonal events, respectively. As shown in **Figure 3B-C**, Cosmic Signature 4 remained as the predominant signature in trunk mutations consistent with previous reports that smoking associated mutational processes play critical roles during early mutagenesis of lung cancers^17,19,33^. On the other hand, the contribution of Signature 4 was significantly reduced (p=0.002) while Cosmic Signature 3 (associated with defect of DNA double-strand break-repair) emerged as the predominant signature for non-trunk mutations (p< 0.0001). Taken together, these results highlight the dynamic nature of mutagenesis at different times during evolution of SCLC and suggest that smoking-associated mutational processes play essential roles during early clonal expansion while subclonal diversification of this cohort of SCLC may be associated with other mutational processes such as DNA repair defect.

### Suppressed TCR repertoire in SCLC

We next performed TCR sequencing in 36 tumor specimens (1 to 3 regions per tumor) and 16 tumor-adjacent lung tissues from patients with adequate DNA remaining. T-cell density, an estimate of the proportion of T cells in a specimen, ranged from 0.11% to 33% with a median of 1.7% (**Figure S5A**). T-cell richness, a measure of T-cell diversity, ranged from 38 to 8,286 unique T-cells (median: 510) per specimen (**Figure S5B)** and T-cell clonality, a metric indicating T-cell expansion and reactivity, ranged from 0.002 to 0.139 (median=0.009) (**Figure S5C**). Density, richness and clonality were positively correlated with each other (Density *vs*. Richness: r=0.87, p<0.0001; Density *vs*. Clonality: r=0.88, p<0.0001; Clonality *vs*. Richness: r=0.97, p<0.0001) (**Figure S5D**). Compared to tumor-adjacent lung tissues (≥2 cm from tumor margin), SCLC tumors demonstrated lower T-cell density, richness and clonality (p=0.0580, p=0.0067, p=0.0166, respectively; **Figure S6A-C**), indicating a suppressed T-cell repertoire in tumor tissues consistent with NSCLC^26^. Of particular interest, all three TCR metrics were significantly lower than those of the 216 localized NSCLCs from PROSPECT cohort (T-cell density 0.014 *vs*. 0.21, p<0.0001 (**Figure S7A**); diversity 510 *vs*. 3,246, p<0.0001 (**Figure S7B**); and T-cell clonality 0.009 *vs*. 0.14, p<0.0001 (**Figure S7C**))^26^.

Additionally, we derived immune scores quantifying the density of immune-cells within tumors by deconvoluting RNA seq data of 81 SCLCs^28^ and compared those to 1,027 NSCLCs from TCGA^33,34^. In line with TCR repertoire findings, the immune scores in SCLCs were significantly lower than NSCLCs (**Figure S8**, p<0.0001). Taken together, these results suggested that SCLC may have more suppressed immune microenvironment than NSCLC.

### Substantial TCR repertoire heterogeneity in SCLC

To gain insights into TCR heterogeneity, we calculated Jaccard index (JI), a metric measuring the proportion of shared T-cell clonotypes between two samples. Substantial TCR heterogeneity was evident across all tumors, with a median JI of 0.05 (0.02 to 0.15) in the 10 SCLCs with multiregional TCR data available (**Figure 4A**), significantly lower than the 11 localized NSCLCs^18^ (median 0.05 in SCLC *vs*. 0.16 in NSCLC, p<0.0001) (**Figure S7D**). Furthermore, 79.9%-97.7% of T-cell clones were restricted to individual tumor regions while only 0.2%-14.6% were identified in all regions within the same tumors **(Figure 4B)**, significantly lower than NSCLC (1.6% to 14.5%, p=0.0048)^18^ demonstrating profound TCR ITH in SCLC even beyond NSCLC.

**Figure 4.**
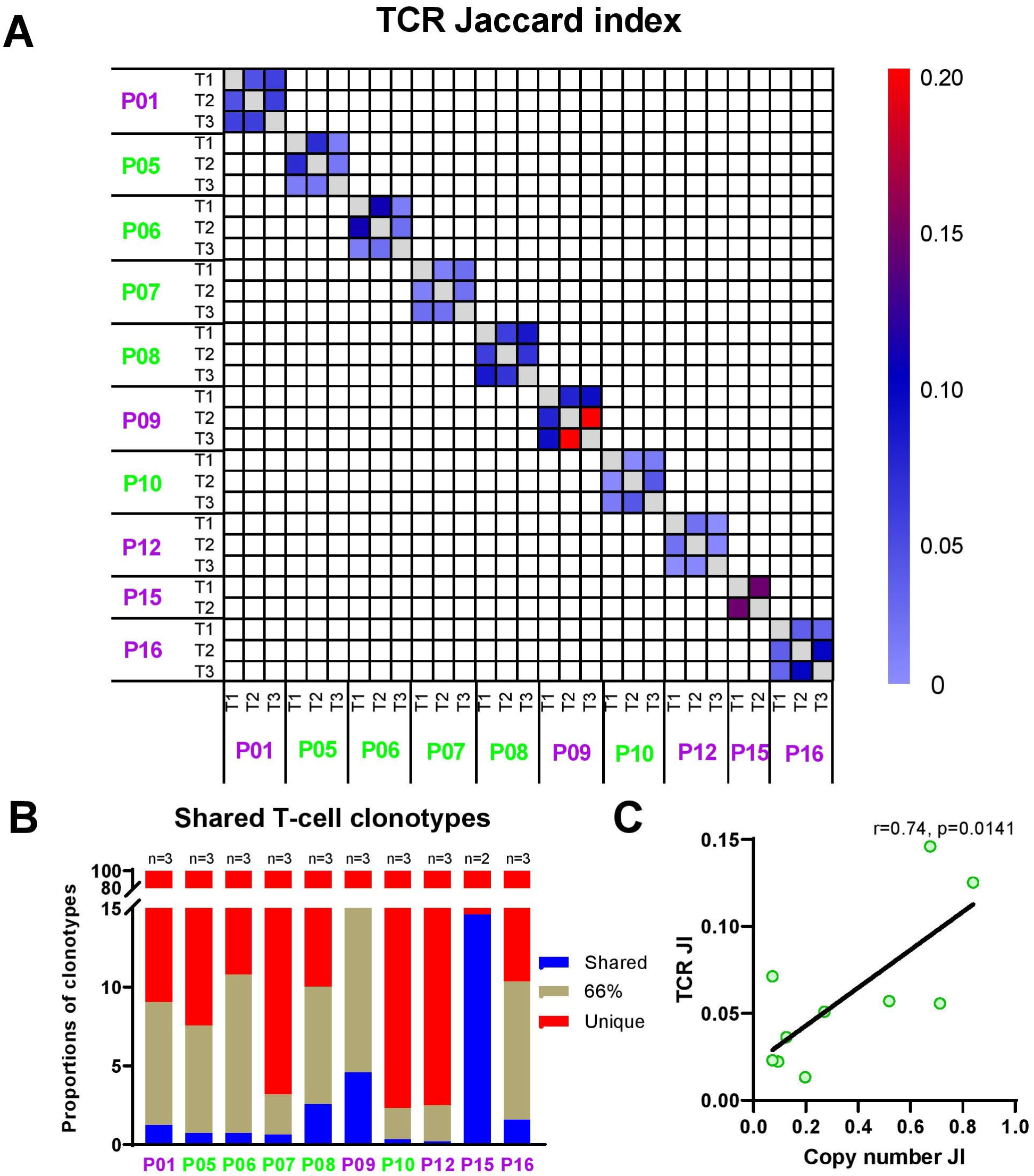
Substantial TCR repertoire intratumor heterogeneity (ITH) in small cell lung cancer. **(A)** Quantification of T cell receptor (TCR) ITH by Jaccard index (JI), a metric representing the proportion of shared T-cell clonotypes between two samples. **(B)** Proportions of T-cell clonotypes detected in all regions (shared, blue), in 2/3 (brown) and restricted to a single region (red) from the same tumors. Patient ID: pink = alive; green = expired. **(C)** Correlations between TCR ITH and TCR ITH by JI.

### High-level and heterogeneous copy number alterations may be the underlying genomic basis for suppressed and heterogeneous TCR repertoire in SCLC

To identify genomic aberrations that could contribute to the suppressive TCR repertoire and ITH in SCLC, we first looked at somatic mutations that play central roles in anti-tumor T cell response by producing non-self proteins that can be recognized by T cells – so called neoantigens^35,36^. We performed *in silico* prediction of HLA-A-, -B-, and -C-presented neoantigens. A median of 78 (26-463) predicted neoantigens (IC_50_< 500 nmol/L) per tumor were detected (**Figure S9A**), which was similar to NSCLCs from the PROSPECT cohort (median: 72/tumor, 2-801, **Figure S9B**, p=0.31). Similar to somatic mutations, 81% (43%-93%) of predicted neoantigens were present across different regions within the same tumors (**Figure S9C**) suggesting the suppressed and heterogeneous TCR repertoire in SCLC was unlikely due to low clonal neoantigen burden.

Next, we explored whether SCLC had higher incidence of loss of heterozygosity (LOH) of human leukocyte antigen (HLA), a potential immune evasion mechanism in cancer ^37,38^. Evidence of HLA LOH was revealed in 9 of 19 SCLCs, higher than NSCLCs from the PROSPECT cohort, but the difference did not reach statistical difference (9/19 vs. 60/216, p=0.11) suggesting that HLA LOH is a common mechanism underlying immune evasion in both SCLC and NSCLC, but may not be the main determinant of more suppressed TCR repertoire in SCLC *versus* NSCLC.

As a higher chromosomal copy number aberration (CNA) burden has been reported to associate with immunosuppression in multiple cancer types^39,40^, we next assessed the CNA burden in this cohort of SCLC. A median of 2,180 CNA events per tumor (26 to 7622) were identified from these SCLCs (**Figure S10A**), significantly higher than 622 per tumor (range: 0-7741) in NSCLCs from PROSPECT cohort (p<0.0001) (**Figure S10B**)^26^. Additionally, in this cohort of SCLC tumors, CNA burden was negatively associated with T-cell density, richness and clonality (r=−0.4, p=0.0157; r=−0.36, p=0.0317; r=−0.33, p=0.0484; respectively) (**Figure S11A-C**). Furthermore, CNA JI, a surrogate for CNA ITH was positively associated with TCR JI (r=0.74, p=0.0141) (**Figure 4C)**. Taken together, these data suggest that higher CNA burden and higher level of CNA ITH could be important genomic basis for profoundly suppressed and heterogeneous TCR ITH in these SCLCs.

### Genomic and TCR ITH were associated with survival of SCLC

With small sample size fully acknowledged, we attempted to assess whether the genomic and T cell features impact clinical outcome. We focused on overall survival since recurrence status was unavailable for some patients. With a median of 45 months of postsurgical follow up, 9 patients have expired with a median of OS of 45 months, comparable to previous reports^41-44^. Interestingly, higher TMB was associated with significantly longer OS (**Figure 5A**, HR=0.13, p=0.0281), consistent with previous reports in NSCLC^45^. Conversely, higher CNA burden was associated with significantly shorter OS (**Figure 5B**, HR=13.8, p=0.0033), while significantly longer OS was observed in patients with low level of copy number ITH (high CNA JI, **Figure 5C**, HR=4.21E-10, p=0.0019). No TCR parameter (T-cell density, richness and clonality) was associated with OS (**Figure S12A-C**), however, patients with a more homogenous TCR repertoire (higher TCR JI) exhibited significantly longer OS (**Figure 5D**, HR=0.16, p=0.0496).

**Figure 5.**
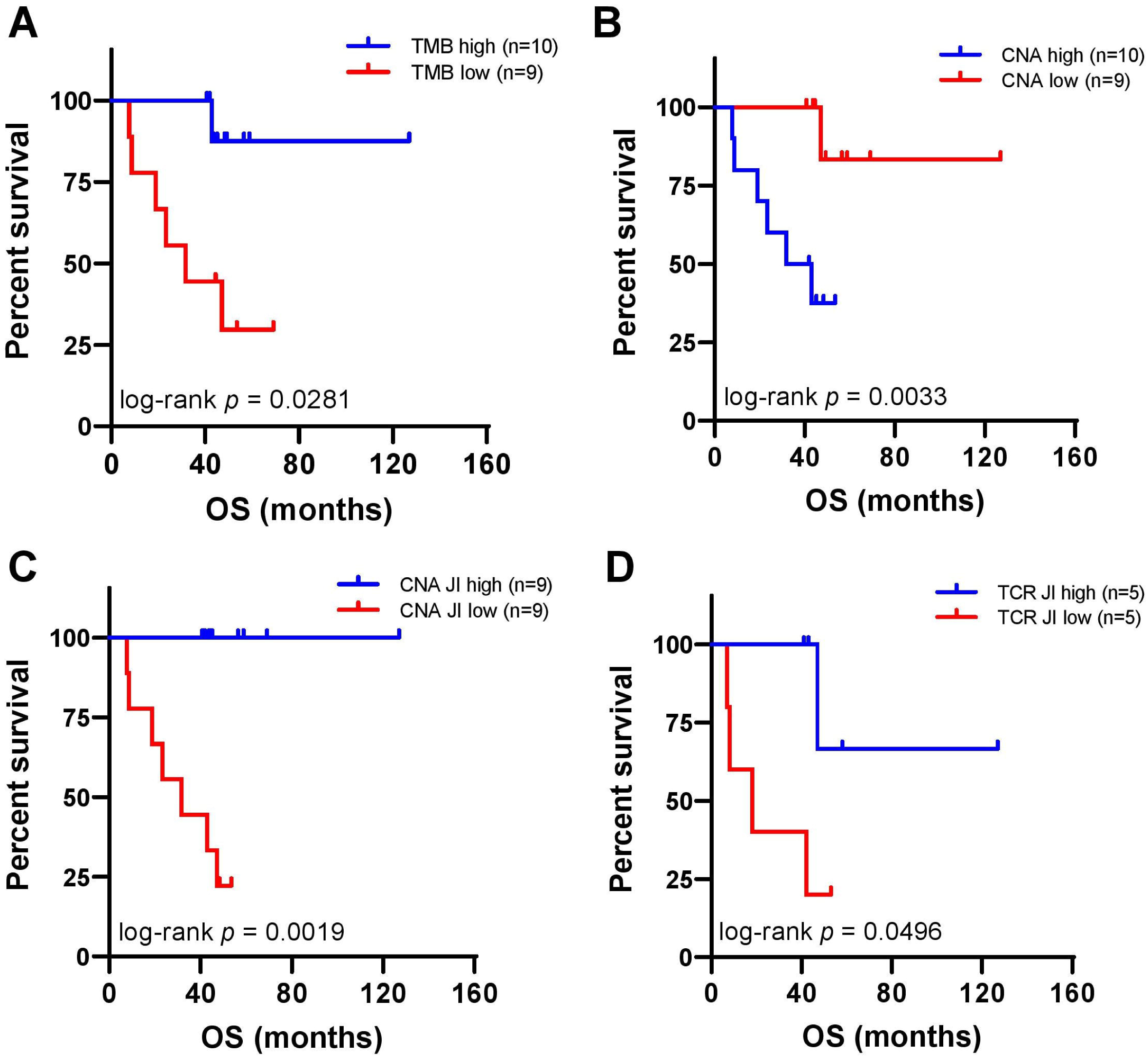
Association of overall survival (OS) with genomic and TCR landscape. **(A)** OS is longer in patients with higher (above median, blue) TMB than patients with lower (below median, red) TMB. **(B)** OS is shorter in patients with higher (above median, blue) CNA burden than patients with lower (below median, red) CNA burden. **(C)** OS is longer in patients with less CNA ITH (higher CNA JI, blue) than patients with higher level of CNA ITH (lower CNA JI, red). **(D)** OS is longer in patients with more homogenous TCR repertoire (higher above median TCR JI, blue) than patients with more heterogeneous TCR repertoire (lower below median TCR JI, red).

## Discussion

Evolutionary theory suggests populations of high genetic variation have survival advantages^46^. Similarly, tumors of complex ITH may be difficult to eradicate. Higher level of molecular ITH has been demonstrated to associate with inferior outcome of cancer patients^12,16,17^. In SCLC, however, although pioneering studies have revealed some pivotal molecular features^27-30,47^, the genomic ITH architecture has not been defined, primarily due to the lack of adequate tumor specimens for multiregional profiling. Because SCLC is sensitive to initial treatment but nearly all patients experience relapse with refractory disease, it has been speculated that SCLC may have profound mutational ITH, where cancer cells highly resistant to chemotherapy/radiotherapy hide in the treatment-naïve SCLC tumors as minor subclones that give rise to relapse^7,11^. Surprisingly, all SCLCs in the current study demonstrated homogeneous mutational ITH with the majority of mutations present in all regions within the same tumors (**Figure 1 and 2A**) and a median of 92.8% of mutations being clonal in each tumor specimen (**Figure 2B**). Additionally, previous work from Wagner and colleagues has demonstrated striking similarity of the mutational landscape between primary and relapsed SCLC^47^. Taken together, these data indicate that complex mutational ITH and selection of chemo-/radio-resistant minor subclones may not be the main mechanisms underlying therapeutic resistance in SCLC.

Cancer evolution with or without treatment may be shaped by the dynamic interaction between cancer cells and host factors, particularly through immune surveillance^48^. Our study delineates for the first time, the TCR repertoire of SCLC and demonstrates a suppressed T-cell repertoire in SCLC. All TCR attributes were extremely low quantitatively (density) and qualitatively (richness and clonality), compared to not only matched normal lung tissues (**Figure S6**) but also compared to NSCLC tumors (**Figure S7A-C**). Similarly, comparing a previously published large SCLC cohort (n=81)^28^ to TCGA NSCLC cohorts (n=1,027) also revealed more suppressed immune contexture in SCLC than NSCLC (**Figure S8**).

In addition to the suppressed TCR repertoire, SCLC also demonstrated extremely heterogeneous TCR repertoire with only 0.2%-14.6% of all T cells identified across all tumor regions within the same tumors. TCR ITH was even more pronounced than that in NSCLC (**Figure S7D**)^18^, which may further impair the efficacy of anti-tumor immune response. Interestingly, even with such a small sample size, higher TCR JI indicating less TCR ITH was associated with better survival in these SCLC patients (**Figure 5B**) indicating the potential clinical impact of TCR ITH. SCLC is among the cancers with high TMB^49^ and our study also demonstrated homogenous mutational landscape, both of which have been reported to associate with benefit from ICB^50^. However, compared to NSCLC and other tumor types, fewer SCLC patients benefit from ICB^51^. The suppressed and heterogeneous TCR repertoire may be one potential reason underlying suboptimal response to immunotherapy.

As the TCR repertoire attributes in this cohort of SCLC were significantly suppressed compared to NSCLC from PROSPECT cohort (**Figure S7A-C**), we compared the genomic landscape of tumors of these two cohorts to understand the potential genomic bases for the more suppressed TCR repertoire in SCLC. These analyses revealed significantly higher CNA burden in SCLC (**Figure S10B**). Moreover, the CNA burden was negatively associated with both T cell quantity (density) and quality (richness and clonality) (**Figure S11**) and CNA ITH was positively associated with TCR ITH (**Figure 4C**) in this cohort of SCLC. These results suggest that high CNA burden and high level of CNA ITH may be one of the underlying genomic bases for the suppressed and heterogeneous TCR repertoire in this cohort of SCLC.

High CNA burden has been reported to correlate with immunosuppressive microenvironment and inferior benefit from ICB across different cancer types^39,52,53^. The mechanisms underlying the association between high CNA burden and immunosuppression are not well understood. Several hypotheses have been proposed such as relatively low neoantigen concentration due to protein imbalance leading to impaired cancer cell signals in tumors^39,52^. From a therapeutic standpoint, the significantly higher CNA burden suggests targeting CNA could be a potential effective strategy for treating SCLC. Although CNA can potentially lead to gene dosage effects that could promote tumor growth and provide the immune evasive advantage for cancer proliferation^39,52,53^, excessive CNA beyond a certain level could be lethal to cancer cells^54,55^. Genes and pathways involved in CNA (e.g. spindle assembly checkpoint, supernumerary centrosome clustering, Aurora kinase family members, etc.) have been exploited as candidate therapeutic targets for different cancer types including SCLC^56^. Unfortunately, none of these agents has shown substantial efficacy to make the way to clinical practice in treating SCLC although anti-tumor activities have been observed from several agents of this class^54,56,57^. One plausible explanation is the profound CNA ITH in SCLC as observed in the current study where different cancer cells may have vastly different CNA profiles. As such, these CNA promoting agents could kill cancer cells with excessive CNA while spare cancer cells with less CNA leading to therapeutic failure. Moreover, increasing CNA potentially turns formerly CNA-low cancer cells into relatively CNA-high cells starting the cycle again that further suppressing host anti-tumor immune response. This is a similar quandary with inhibiting DNA damage response (DDR) pathway where deficient DDR pathways could increase DNA-damaging chemo-/radio-therapy sensitivity but conversely promote tumorigenesis^58,59^. Therefore, in order to effectively eliminate heterogeneous cancer cells with different CNA profiles, CNA targeting agents may be combined with ICB, which has already been tested in treating SCLC in both preclinical murine models ^60^ and clinical trials (NCT03041311)^61^.

To the best of our knowledge, the current study is the first study on genomic and TCR ITH of SCLC. Our study was limited by the small sample size due to the scarcity of resected SCLC specimens. However, WES and TCR data from multiregional specimens made it valuable to the field. In summary, we demonstrate that despite a homogeneous mutational landscape, SCLC exhibits a suppressed and heterogeneous TCR repertoire that could lead to ineffective anti-tumor immune surveillance, which could be one potential molecular mechanism underlying high recurrence rate and suboptimal response to immunotherapy in SCLC. Our results also suggest that high CNA burden may be one of the underlying reasons for the suppressed T-cell repertoire, therefore a potential therapeutic target to improve the efficacy of immune checkpoint blockade in patients with SCLC.

## Methods

### Patients

A total of 19 patients with lymph node negative LD SCLC, who underwent surgical resection at Zhejiang Cancer Hospital, Hangzhou, China from 2010 to 2015 were enrolled. With a median of 45 months of postsurgical follow up, 7 patients have relapsed and deceased, 10 patients were still alive with no evidence of recurrence and 2 patients deceased with unknown recurrence status. The median survival of this cohort was 45 months. The study was approved by the Institutional Review Boards (IRB) at MD Anderson Cancer Center and Zhejiang Cancer Hospital.

### Sample processing and DNA extraction

Hematoxylin and eosin slides from each tumor were reviewed by experienced lung cancer pathologists to confirm the diagnosis, assess necrosis, tumor purity and cell viability. Manual macro-dissection was conducted to enrich malignant cells. DNA was extracted using the AllPrep® DNA/RNA FFPE Kit (Qiagen, Hilden, Germany) from 50 spatially separated tumor regions (3 regions per tumor from 13 patients, 2 regions per tumor from 5 patients and 1 tumor piece from one patient) and paired matched adjacent normal lung (≥2 cm from tumor margin, morphologically negative for malignant cells assessed by two lung cancer pathologists independently) as previously described^62^.

### Whole exome sequencing

WES was performed using the Illumina protocol in MD Anderson. Exome capture was performed on 500ng of genomic DNA per sample based on KAPA library prep (Kapa Biosystems) using the Agilent SureSelect Human All Exon V4 kit according to the manufacturer’s instructions and paired-end multiplex sequencing of samples was performed on the Illumina HiSeq 2000 sequencing platform. The average sequencing depth was 180x for tumor DNA (ranging from 64x to 224x), 161x for germline DNA (ranging from 96x to 194x).

### Mutation calling

The BWA aligner (bwa-0.7.5a) was applied to map the raw reads to the human hg19 reference genome (UCSC genome browser: genome.ucsc.edu). The Picard (v1.112, http://broadinstitute.github.io/picard/) ‘‘MarkDuplicates’’ module was applied to mark the duplicate reads. Then the ‘‘IndelRealigner’’ and ‘‘BaseRecalibrator’’ modules of the Genome Analysis Toolkit were applied to perform indel realignment and base quality recalibration. Mutect (v1.1.4) ^63^ was applied identify somatic single nucleotide variants (SNVs) and small insertions/deletions. To ensure high-quality mutation calls, the following filtering criteria were applied: 1) sequencing depth ≥20× in tumor DNA and ≥10×Lin germline DNA; and 2) variant alleleve frequency (VAF) ≥ 0.02 in tumor DNA and < 0.01 in germline DNA; and 3) the total number of reads supporting the variant calls is ≥4; and 4) variant frequency is < 0.01 in ESP6500, 1000 genome and EXAC databases; and 5) LOD score >18 (MuTect default is 6.3). We kept the mutations that passed all filtering criteria except LOD score < 18 only if the identical mutations were present with LOD score >=18 in other regions within the same tumors.

Cancer gene mutations were defined as identical oncogene mutations previously reported; stop gains and frameshift of tumor suppressor genes; other non-synonymous mutations with Combined Annotation Dependent Depletion (ACDD) score>20^64^.

### Clonal and subclonal analysis

Tumor contents and major/minor copy number changes were estimated by Sequenza (v2.1.2).^65^ The cancer cell fraction (CCF) and mutant allele copy number for each SNV was inferred using Pyclone 12.3^31^. In brief, PyClone implements a Dirichlet process clustering model that simultaneously estimates the distribution of the cellular prevalence for each mutation. Copy numbers of somatic mutations were inferred by integrating integer copy numbers determined by Sequenza on single sample basis. The outputs were cellular prevalence value distributions per SNV estimated from Markov-chain Monte Carlo (MCMC) sampling. The median value of the MCMC sampling-derived distribution was used as a representative cellular prevalence for each mutation. A given mutation was classified as “clonal” if the 95% confidence interval of CCF overlapped 1 and “subclonal” otherwise.

### Phylogenetic analysis

Mutation profiles were converted into binary format with 1 being mutated and 0 otherwise. Ancestors were germ line DNA assuming with no mutations. Multistate discrete-characters Wagner parsimony method in PHYLIP (Phylogeny Inference Package) was used to generate phylogenic tree^66^.

### Mutational signature analysis

The R package “DeconstructSigs” package ^67^ was applied to estimate the proportions of 30 COSMIC mutational signatures (http://cancer.sanger.ac.uk/cosmic/signatures).

### Somatic copy number analysis

Somatic copy number analysis were performed applying CNVkit (v0.9.6)^68^, through which both the targeted reads and the nonspecifically captured off-target reads were used to infer copy number evenly across the genome, and DNA segmentation of log2 ratios in the tumor samples were calculated, then segment data were processed using the “CNTools” package to generate segmented DNA copy number profile at gene level by assigning segment means to the genes within the chromosome segments for each sample. Genes with mean segment more than 0.6 was defined as copy number gain and less than −0.6 was defined as copy number loss. Copy number gain and loss burden were defined as the number of genes located in the segments with copy number gains and losses.

### Neoantigen prediction

WES data were reviewed for non-synonymous exonic mutations. The binding affinity with patient-restricted MHC Class I molecules of all possible 9- and 10-mer peptides was evaluated with the NetMHC3.4 algorithm based on patient HLA-A, HLA-B, and HLA-C alleles^69-71^. Candidate peptides were considered HLA binders when IC50<500 nM.

### TCRβ sequencing and comparison parameters

Immunosequencing of the CDR3 regions of human TCRβ chains was performed using ImmunoSeq (Adaptive Biotechnologies, hsTCRβ Kit)^18,26^. T-cell density was calculated by normalizing TCR-β template counts to the total amount of DNA for TCR sequencing, where the amount of total DNA was determined by PCR-amplification and sequencing of housekeeping genes expected to be present in all nucleated cells. T-cell richness is calculated using the unique rearrangements. T-cell clonality is defined as 1-Peilou’s evenness and is calculated on productive rearrangements as previously described^18,26^. Jaccard index (JI) was calculated by the number of rearrangements shared/sum of total number of rearrangements between any two specimens.

### Human leukocyte antigen loss of heterozygosity analysis

For Human Leukocyte Antigen Loss Of Heterozygosity (HLA LOH) analysis, we first performed HLA typing using PHLAT^72^. For each patient, we merged tumor and normal BAM files and inferred 4-digit HLA types for the major class I HLA genes (HLA-A, HLA-B and HLA-C). To evaluate HLA loss, we used a computational tool, LOHHLA ^73^ using purity and ploidy information estimated by Sequenza^74^. Sample as being subject to HLA loss was defined when any of the two alleles of HLA-A, HLA-B or HLA-C showed a copy number□<□0.5 with a paired Student’s *t* test *p*□<□0.01.

### Analysis of published data

RNA sequencing data from 81 SCLCs^28^ and 1,027 NSCLCs from TCGA^33,34^ were downloaded. Immune scores were calculated by taking the average of normalized expression levels of genes including cytolytic markers, HLA molecules, IFN-γ pathway, chemokines and adhesion molecules as previously described^75^.

### Statistical Analysis

Graphs were generated with GraphPad Prism 8.0 (La Jolla, CA). Pearson’s correlations were calculated to assess association between 2 continuous variables. Wilcoxon signed-rank test was applied to compare paired TCR metrics. Mann-Whitney test was used to compare differences between two independent groups. Log-rank test was used for survival analysis.

## Acknowledgements

Barbanti Small Cell Lung Cancer Award, Conquer Cancer Foundation ASCO Young Investigator Award, MD Anderson Physician Scientist Award, University Cancer Foundation Sister Institution Network Fund, Cancer Prevention * Research Institute of Texas (CPRIT) Multiple Investigator Award, TJ Martell Foundation, NIH/NCI R01-CA207295, NIH/NCI U01-CA213273, Department of Defense (LC170171), National Natural Science Foundation of China (grant No. 81672972); Major Program of Provincial Ministry of Health Science Foundation (grant No. WKJ-ZJ-1701).

## Author contributions

M.C. and J.Z. conceived the study. R.C., J.L. and Jiexin Z. led the data analysis. J.Y., J.F, H.P., CW.C. and J.F. led the pathological assessment, multi-region sample preparation and DNA extraction. J.Y, Y.C. and X.H. collected resected specimens and clinical data. L.L and C.G. performed DNA preparation and whole-exome sequencing. X.S. and Jianhua Z. performed sequencing raw data processing. J.L., W.L. and X.H. performed downstream bioinformatics analyses. R.C., S.M.H, J.G., C.B., E.R.P., C.G., Robert C., D.G., J.H., W.W., B.G., I.W., P.A.F., R.K.T., A.R., L.A.B. and J.Z. interpreted the data for clinical and pathological correlation. L.D., Q.W., J.W., and JJ.L. performed statistical analyses. R.C., A.R., C.G. and J.Z. wrote the paper. All authors edited the manuscript.

