## Supplemental figures and Table S1 for "Genomic and TCR Repertoire Intratumor Heterogeneity of Small-cell Lung Cancer and its Impact on Survival"

**Supplemental documents**
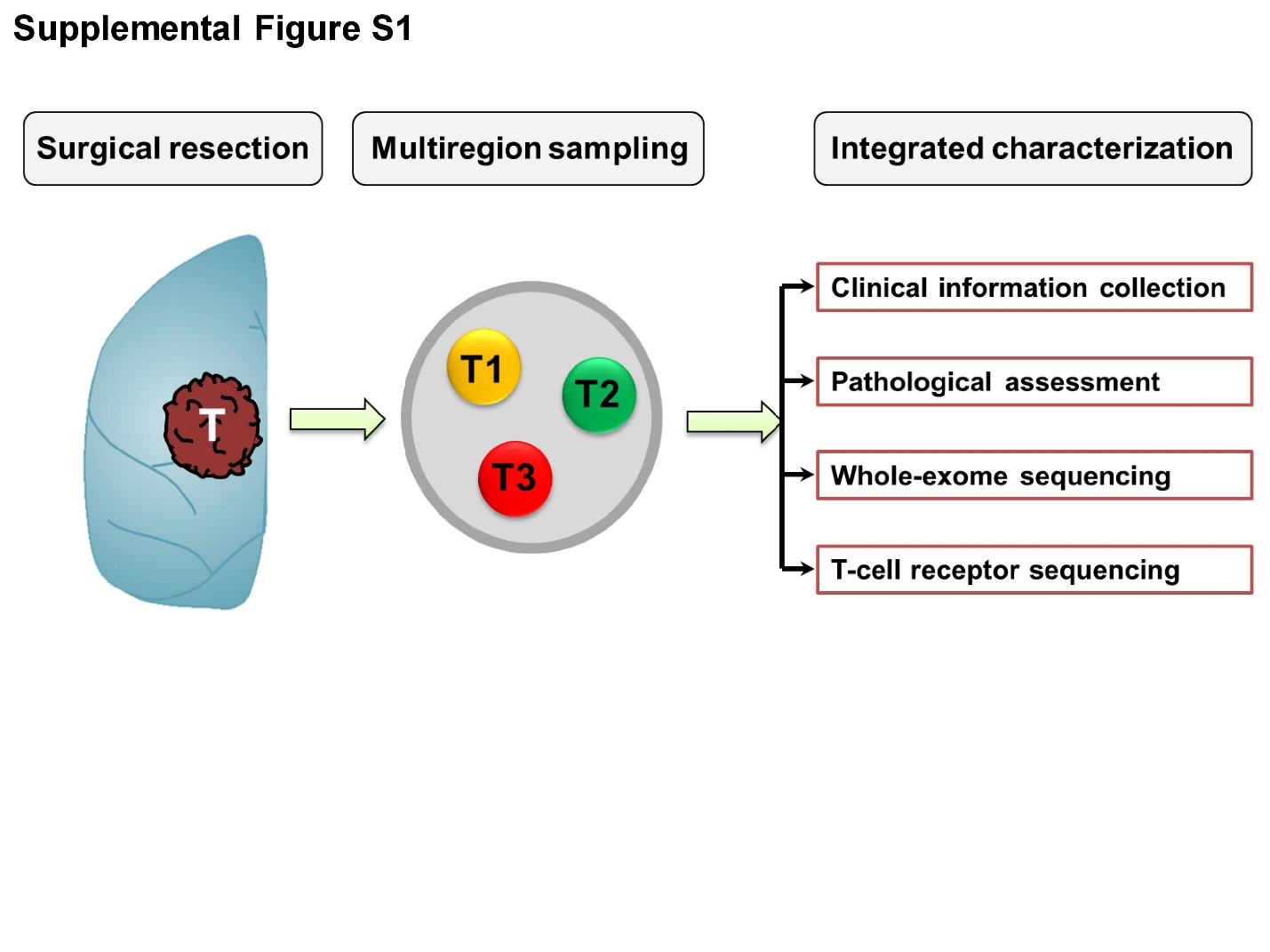

**Supplementary Figure 1.  Study scheme of genomic and T cell receptor (TCR) ITH of SCLC by multiregional sequencing.**

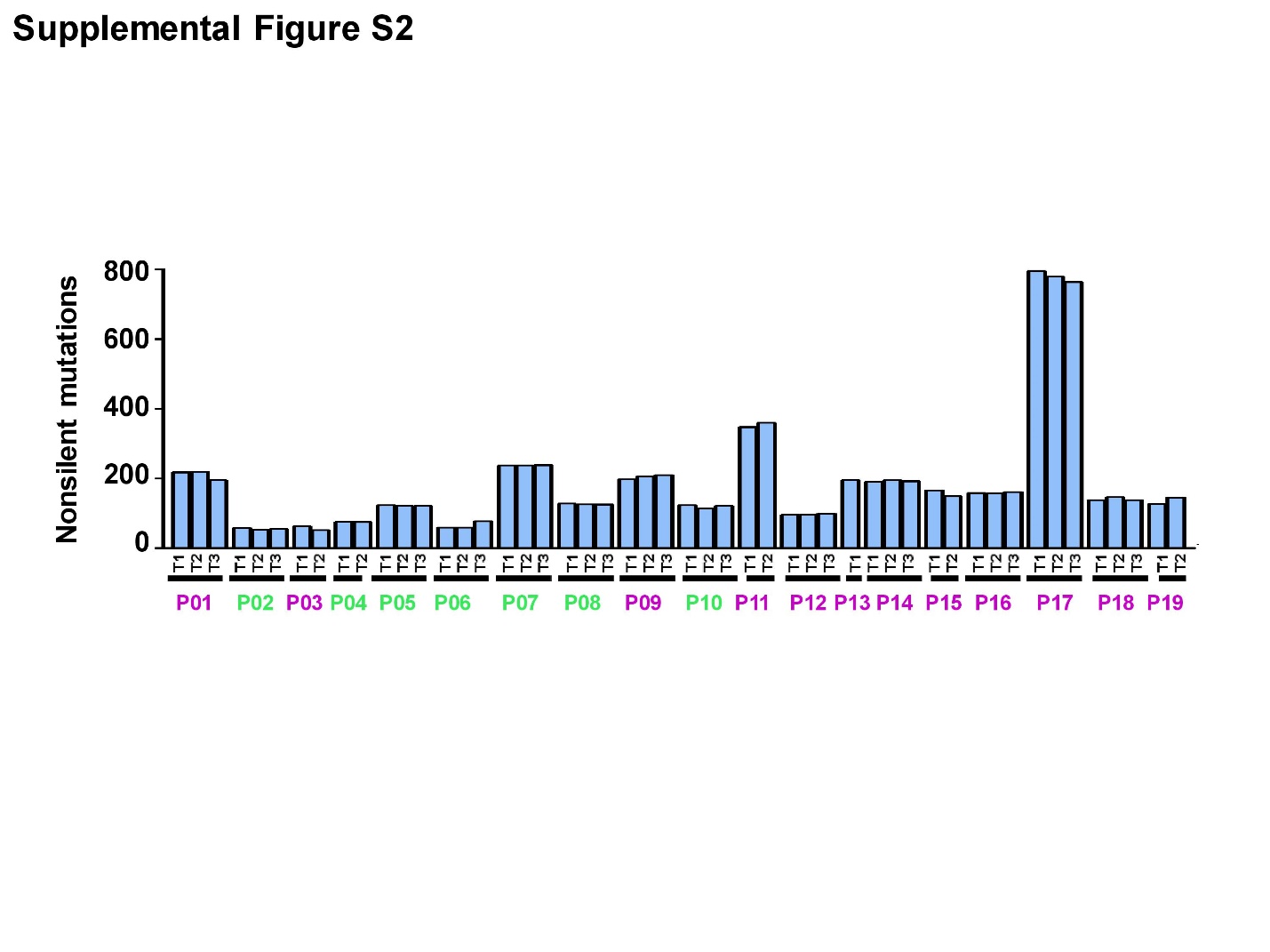

**Supplementary Figure 2. Total number of non-silent mutations identified in 50 SCLC specimens.** Patient ID: pink = alive; green = expired.

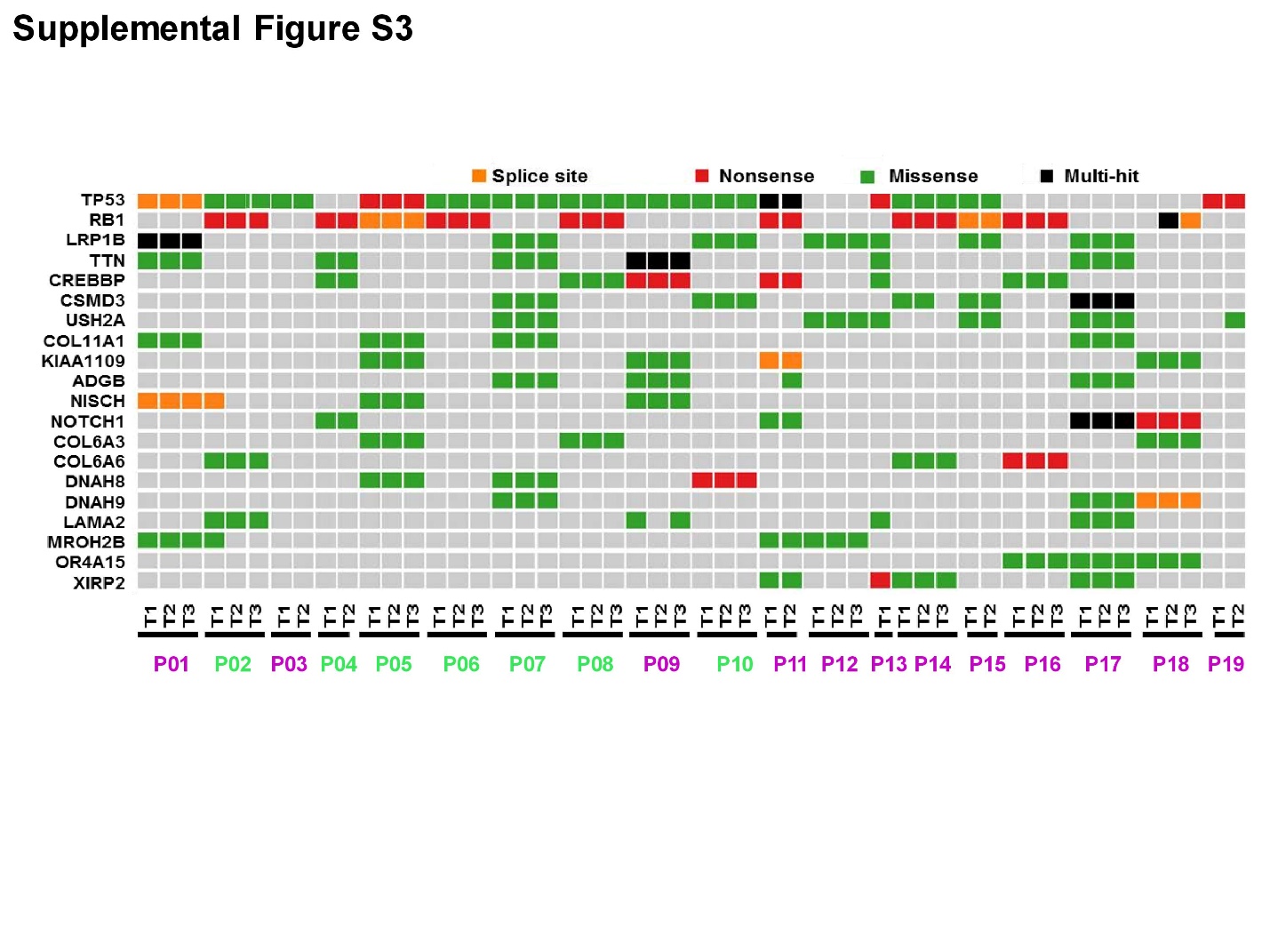

**Supplementary Figure 3. Cancer gene mutations in 50 SCLC specimens.** Cancer gene mutations included identical mutations previously reported in oncogene, stop gain or frameshift mutations in tumor suppressor genes and other non-silent mutations with Combined Annotation Dependent Depletion (ACDD) score>20. Patient ID: pink = alive; green = expired.

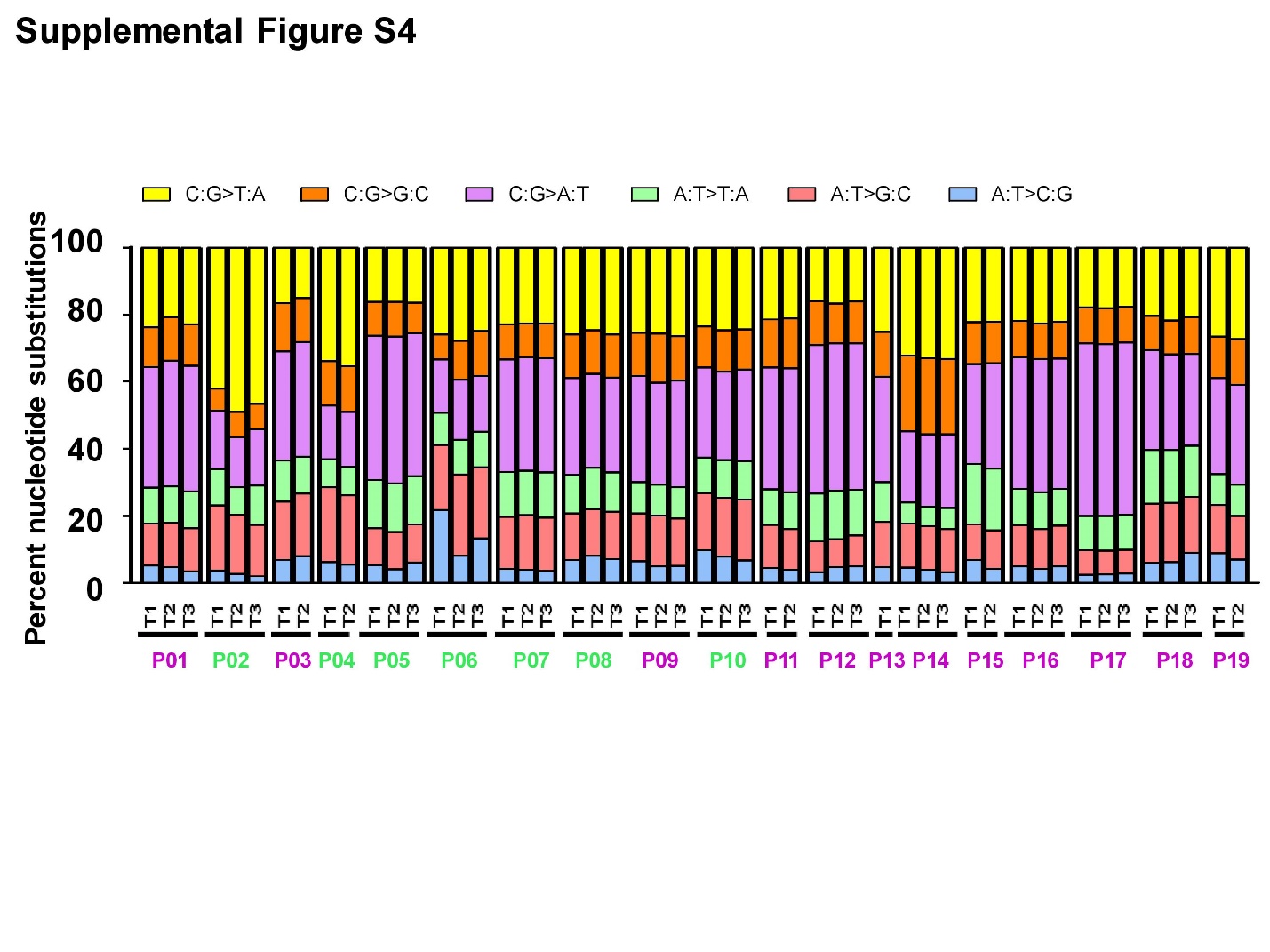

**Supplementary Figure 4. Mutation spectrum in 50 SCLC specimens.** Patient ID: pink = alive; green = expired.

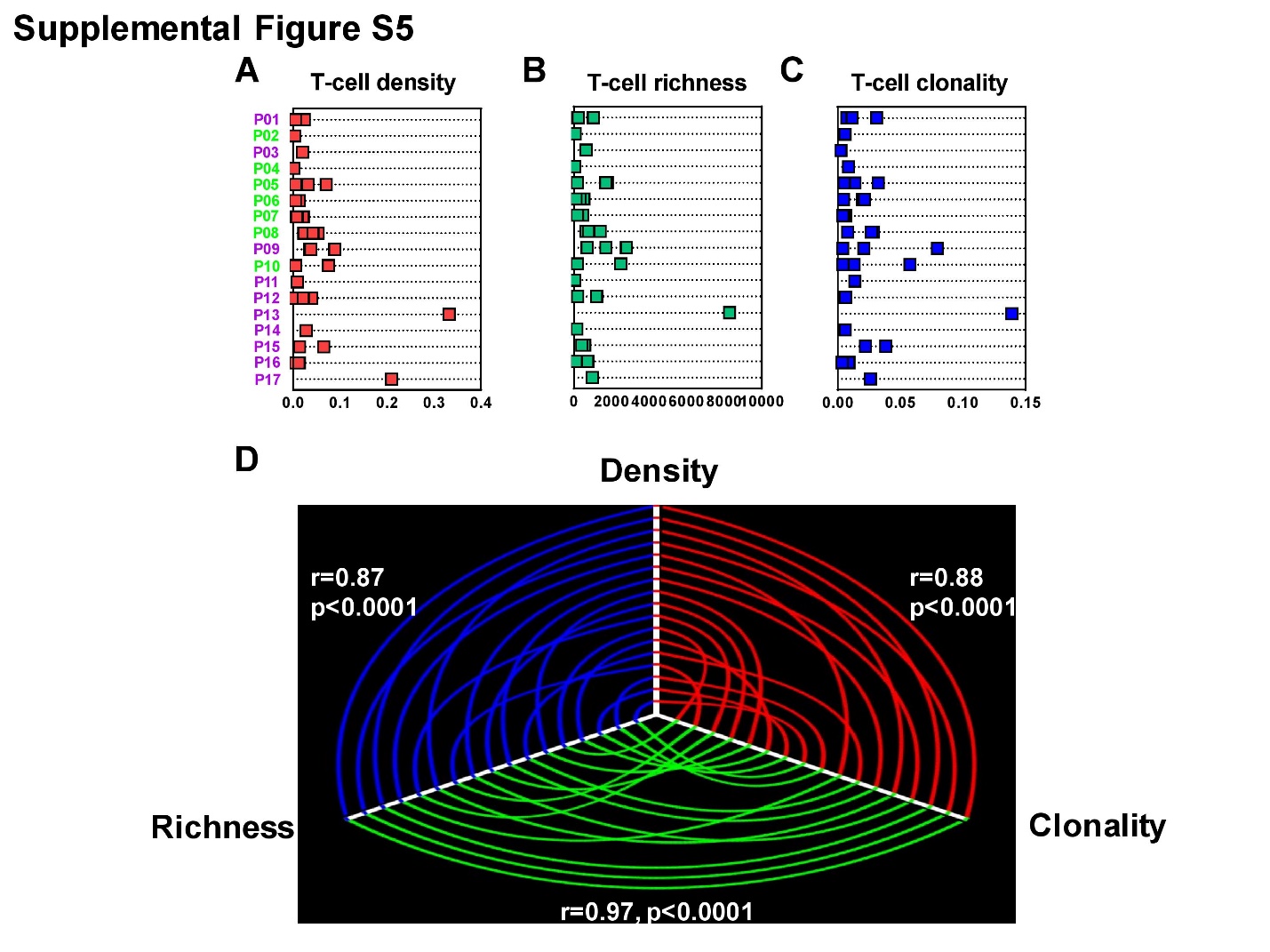

**Supplementary Figure 5. T cell receptor (TCR) repertoire landscape of SCLC. (A)** T-cell density (red) – an estimate of the proportion of T cells in a specimen. **(B)** T-cell richness (green) - a measure of T-cell diversity. **(C)** T-cell clonality (blue) - a metric indicating T-cell expansion and reactivity. **(D)** TCR metrics were positively inter-correlated. Patient ID: pink = alive; green = expired.

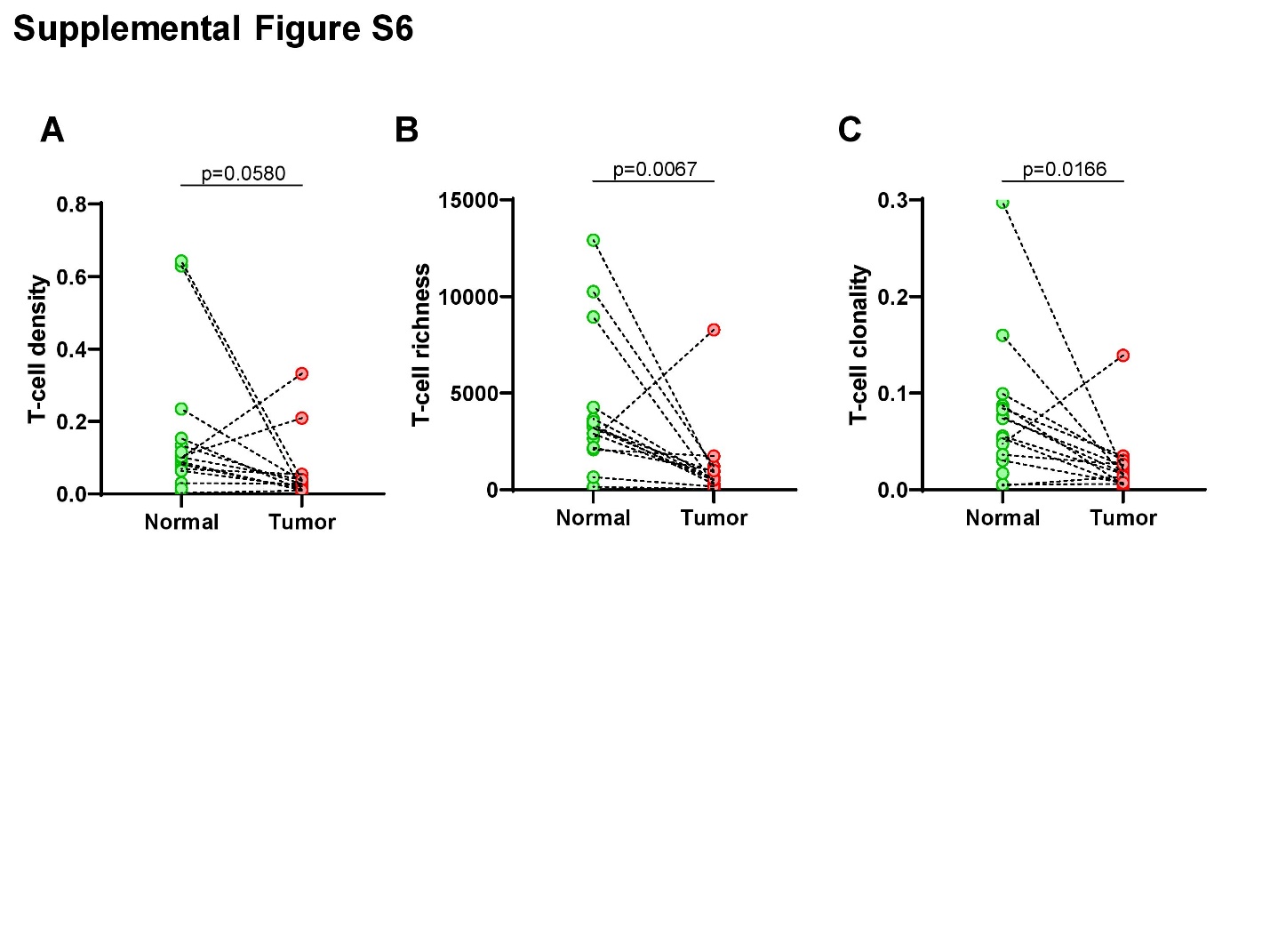

**Supplementary Figure 6. Comparison of TCR metrics between SCLCs tumors and normal lung tissues.** T-cell **(A)** density, **(B)** richness and **(C)** clonality in tumors (red) *versus* normal tissues (green).

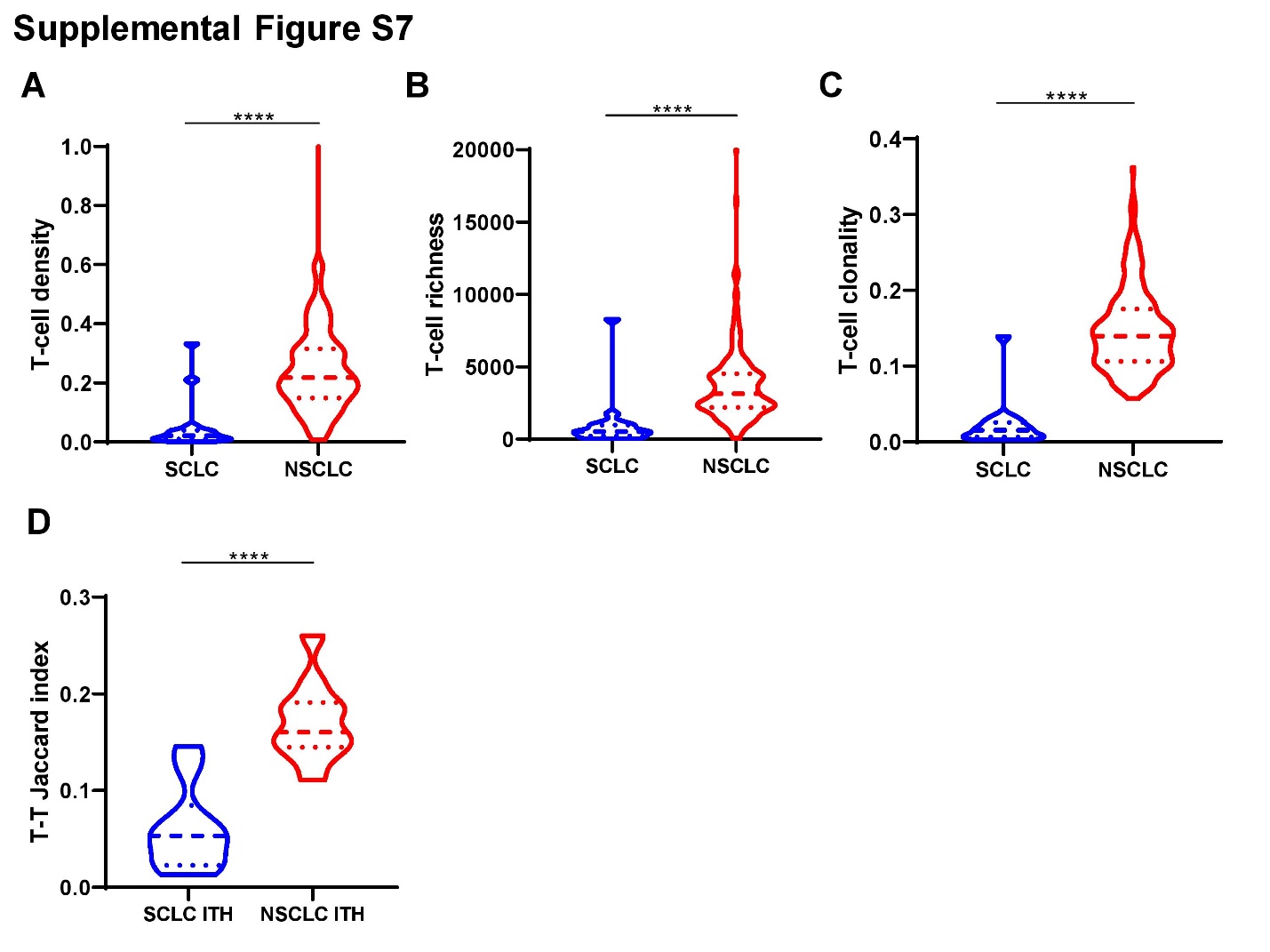

**Supplementary Figure 7. Comparison of TCR metrics between SCLCs and NSCLC tumors from PROSPECT cohort. (A)** T-cell density - an estimate of the proportion of T cells in a specimen. (**B)** T-cell richness - a measure of T-cell diversity and **(C)** T-cell clonality - a metric indicating T-cell expansion and reactivity in SCLC (blue) *versus* NSCLC (red). **(D)** TCR intra-tumor heterogeneity using the average Jaccard index (JI), a metric representing the proportion of shared T-cell clonotypes between two samples in 10 SCLC (blue) *versus* 11 NSCLC tumors (red) with multiregional TCR data.

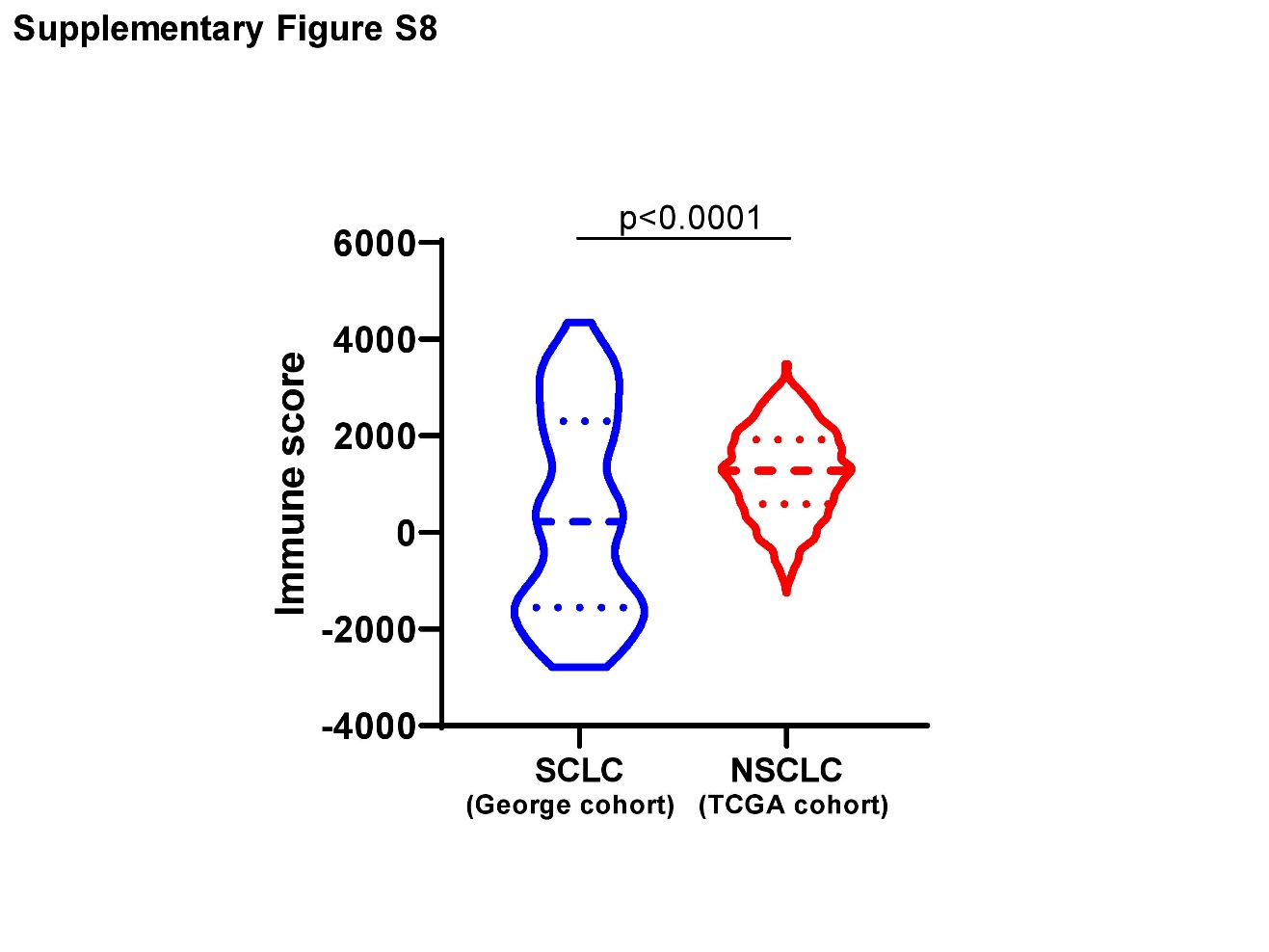

**Supplementary Figure 8. Comparison of immune scores in SCLCs *versus* NSCLCs.** The immune scores were derived from RNA-seq data to quantify the density of immune-cells within the tumors from 81 SCLC tumors (George cohort) *versus* 1,027 NSCLC tumors from TCGA.

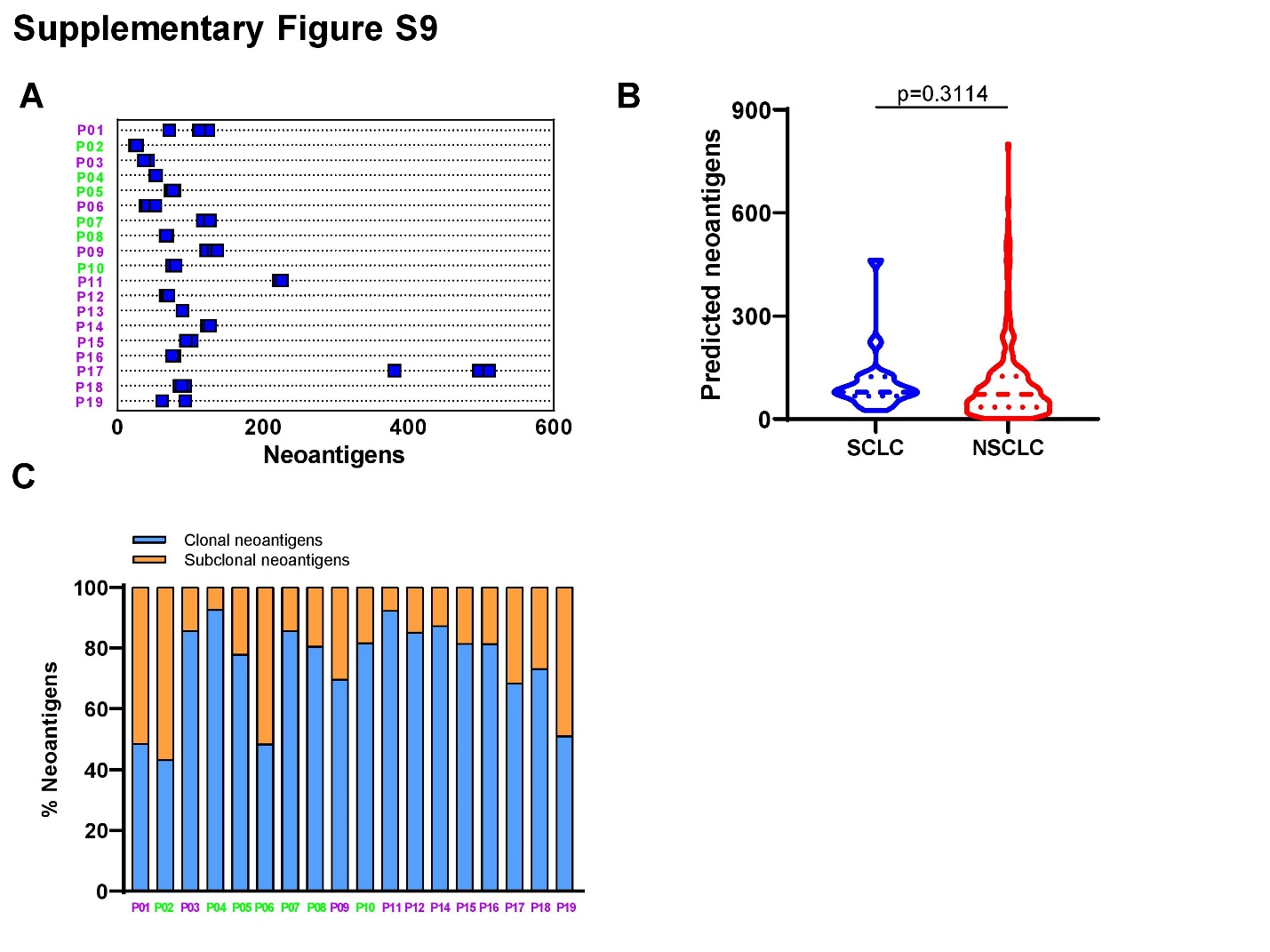

**Supplementary Figure 9. Predicted neoantigen of SCLCs *versus* NSCLCs. (A)** Predicted neoantigens in SCLCs. **(B)** Predicted neoantigen burden in SCLCs *versus* NSCLCs from PROSPECT cohort. **(C)** Proportion of predicted neoantigens associated with clonal (blue) *versus* subclonal (orange) mutations in 18 SCLCs with multiregional exome sequencing data available. Patient ID: pink = alive; green = expired.

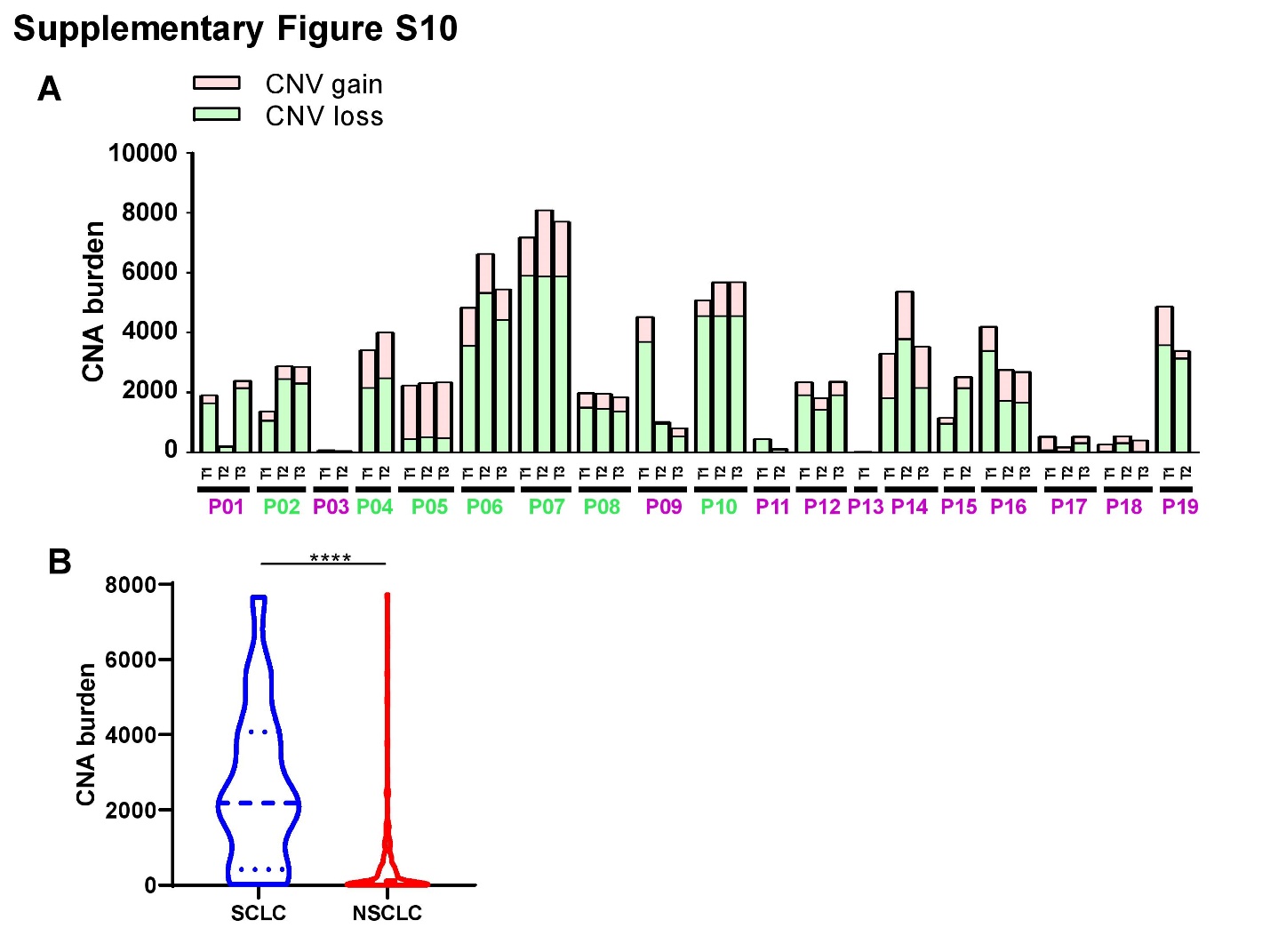

**Supplementary Figure 10. Copy number alterations (CNA) in SCLCs *versus* NSCLCs. (A)** Copy number gain burden (number of genes in the chromosomal segments with tumor/normal log2 ratio ≥ 0.6, peach) and loss burden (number of genes in the chromosomal segments with tumor/normal log2 ratio ≤ -0.6, green) in 50 SCLC tumors. **(B)** CNA burden (copy number gain burden + copy number loss burden) in SCLC is significantly higher than NSCLCs from PROSPECT cohort.

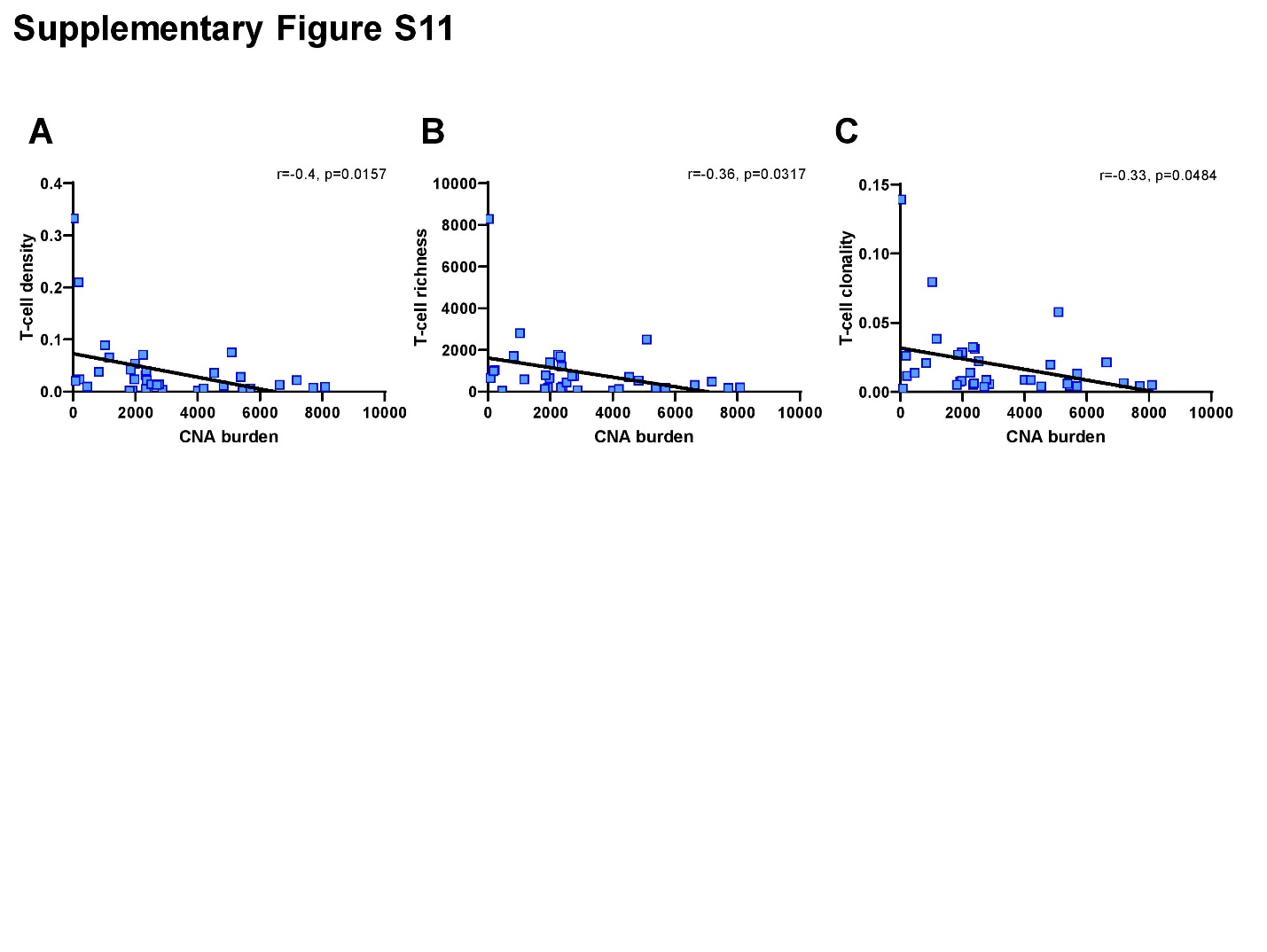

**Supplementary Figure 11.** **Correlations between CNA burden with TCR metrics in SCLCs.** Correlations of CNA burden with **(A)** T-cell density - an estimate of the proportion of T cells in a specimen, **(B)** T-cell richness - a measure of T-cell diversity and **(C)** T-cell clonality - a metric indicating T-cell expansion and reactivity. CNA burden is defined as the number of genes in the chromosomal segments with tumor/normal log2 ratio ≥ 0.6 (copy number gain) or ≤ -0.6 (copy number loss).

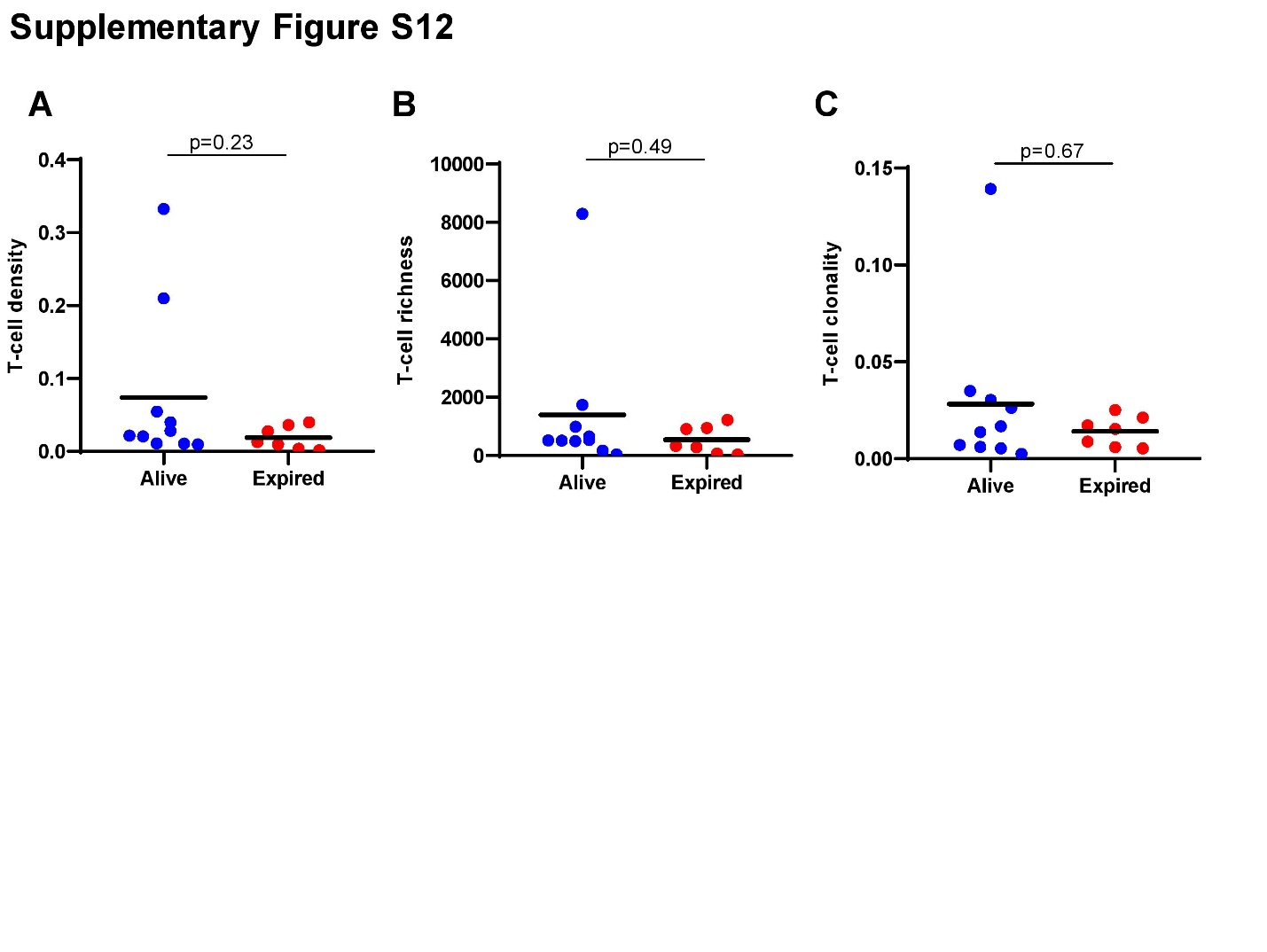

**Supplementary Figure S12. TCR features and vital status of SCLC patients. (A)** T-cell density - an estimate of the proportion of T cells in a specimen, **(B)** T-cell richness - a measure of T-cell diversity and **(C)** T-cell clonality - a metric indicating T-cell expansion and reactivity in SCLC patients who were alive (blue) *versus* expired (red).

**Supplementary Table 1. Clinical characteristics**

| **New ID** | **Gender** | **Age** | **Smoking** | **Recurrence** | **Vital status** | **DFS (Months)** | **OS (Months)** |
| --- | --- | --- | --- | --- | --- | --- | --- |
| **P01** | male | 49 | Yes | No | Expired | 127 | 127 |
| **P02** | female | 62 | No | Yes | Alive | 1 | 23 |
| **P03** | male | 56 | Yes | No | Expired | 45 | 45 |
| **P04** | male | 61 | Yes | Yes | Alive | 10 | 32 |
| **P05** | female | 55 | No | Yes | Alive | 1 | 9 |
| **P06** | female | 74 | Yes | unknown | Alive | 8 | 8 |
| **P07** | male | 70 | Yes | Yes | Alive | 8 | 43 |
| **P08** | male | 61 | Yes | Yes | Alive | 24 | 47 |
| **P09** | male | 65 | Yes | No | Expired | 59 | 59 |
| **P10** | male | 56 | Yes | unknown | Alive | 19 | 19 |
| **P11** | male | 65 | Yes | No | Expired | 56 | 56 |
| **P12** | female | 54 | No | No | Expired | 54 | 54 |
| **P13** | male | 56 | Yes | No | Expired | 49 | 49 |
| **P14** | male | 59 | Yes | Yes | Expired | 14 | 48 |
| **P15** | female | 65 | No | Yes | Expired | 16 | 44 |
| **P16** | male | 56 | Yes | No | Expired | 42 | 42 |
| **P17** | male | 67 | Yes | No | Expired | 41 | 41 |
| **P18** | male | 60 | Yes | No | Expired | 69 | 69 |
| **P19** | male | 61 | Yes | No | Expired | 45 | 45 |
| **DFS:** disease free survival | | |  |  |  |  |  |
| **OS:** overall survival | |  |  |  |  |  |  |

**Supplementary Data 1. Somatic mutations in 50 SCLC tumor specimens**
